# Pharmacological enhancement of slow-wave activity improves cognition and reduces amyloidosis at an early stage in a mouse model of Alzheimer’s disease

**DOI:** 10.1101/2024.10.02.616223

**Authors:** Sedef Kollarik, Dorita Bimbiryte, Aakriti Sethi, Inês Dias, Carlos G. Moreira, Daniela Noain

**Affiliations:** Department of Neurology, University Hospital of Zurich, 8952, Zurich, Switzerland; Neuroscience Centre Zurich (ZNZ), 8057, Zurich, Switzerland; University of Zurich, 8057, Zurich, Switzerland; D-HEST, ETH Zurich, 8092, Zurich, Switzerland; University Center of Competence Sleep & Health Zurich, University of Zurich, 8057, Zurich, Switzerland

## Abstract

Improving sleep in murine Alzheimer’s disease (AD) is associated with reduced brain amyloidosis. However, the window of opportunity for successful sleep-targeted interventions regarding reduction of pathological hallmarks and related cognitive performance remains poorly characterized. Here, we enhanced slow-wave activity (SWA) during sleep via sodium oxybate (SO) oral administration for 2 weeks at early (6 months old) or moderately late (11 months old) disease stages in Tg2576 mice, and evaluated resulting neuropathology and behavioral performance. We observed that cognitive performance of 6 months old Tg2576 mice significantly improved upon SO treatment, whereas no change was observed in 11 months old mice. Histochemical assessment of amyloid plaques demonstrated that SO-treated 11 months old Tg2576 mice had significantly less plaque burden than placebo-treated ones, whereas ELISA of insoluble protein fractions from 6 months old Tg2576 mice’ brains indicated lower Aβ-42/Aβ-40 ratio in SO-treated group vs. placebo-treated controls. Altogether, our results suggest that SWA-dependent reduction of brain amyloidosis leads to alleviated behavioral impairment in Tg2576 mice only if administered early in the disease course, potentially highlighting the key importance of early sleep-based interventions in clinical cohorts.

**Significance statement:** A staggering amount of people worldwide live with AD. Despite extensive efforts from academy and industry, no cure is available to date. Sleep may be a modifiable factor involved in onset, progression and potentially also treatment of AD. However, knowledge regarding important aspects of sleep-targeted treatments is missing, e.g. adequate window of opportunity for their efficacious application, their minimal duration or their effect onto both pathology and cognition. We found that pharmacologically enhanced sleep depth is associated with reduced amyloid neuropathology in AD mice brains at both early- and moderately advanced-stage disease. However, only early-stage disease mice significantly benefited from sleep-enhancing treatment. Our results strongly encourage further mechanistic research and the launch of larger scale preclinical investigations in early AD populations.

## Introduction

Sleep disturbances are believed to contribute to the development of neurodegenerative disorders such as Alzheimer’s disease (AD), as they are highly prevalent in the pre-clinical stages (Ju et al., 2013), and linked to deterioration of disease symptoms. This association between AD and sleep renders the latter a promising target for potential treatment strategies to prevent or delay the onset of AD, and/or to ameliorate the cognitive decline in AD patients (Ju, Lucey, & Holtzman, 2014).

Both animal and human studies demonstrated that extracellular amyloid beta (Aβ) can accumulate in brain regions that regulate sleep/wake patterns, resulting in increased wakefulness at the expense of sleep (Braak & Braak, 1995; Roh et al., 2012). Twenty-four-hour activity rhythm measurements in >300 participants showed that higher sleep fragmentation was associated with higher Aβ burden especially in ApoE4 carriers (Nguyen Ho et al., 2024). Conversely, reversing amyloid plaque deposition by active immunization with synthetic Aβ1–42 in APPswe/PS1δE9 mice prevented sleep disruption (Roh et al., 2012). Reciprocally, impaired sleep affects disease pathology both in AD animals (Kang et al., 2009; Park, Kim, Hwang, & Han, 2023) and patients (Bianchetti et al., 1995; Ooms et al., 2014; Wennberg, Wu, Rosenberg, & Spira, 2017). Subjective sleep impairments (Branger et al., 2016; Spira et al., 2018; Spira et al., 2013; Sprecher et al., 2015; Vicente De Carvalho et al., 2018) and deficits generating non-rapid eye movement sleep (NREMS)’s slow-wave activity (SWA; Mander et al., 2015) anticipated high levels of Aβ burden measured by positron emission tomography. The relationship between self-reported sleep (Kaprio & Koskenvuo, 2002) and 15-26 years post-survey cognitive outcomes in more than 2000 people ≥ 65 years old indicated that especially longstanding short sleep and poor sleep quality are deleterious for cognitive performance (Virta et al., 2013). In a cross-sectional study with asymptomatic AD patients, researchers observed that longer sleep duration was connected with lower Aβ load in regions of the brain characterized by early deposition (Aslanyan et al., 2023), suggesting strategies to improve sleep quality could potentially help delay the onset of cognitive symptoms associated with AD pathology. In fact, midlife sleep length and quality are associated with late life cognitive function. A community-based study also showed that high sleep fragmentation is associated with higher risk of developing AD in a 6-years follow-up period (Lim, Kowgier, Yu, Buchman, & Bennett, 2013). Abnormal expressions of circadian clock genes were observed in mice following chronic sleep deprivation, with these changes being more pronounced in AD mice compared to WT controls (Niu et al., 2022), indicating that sleep disturbances may have a greater impact on individuals with AD.

Notably, studies suggest that restoration of sleep may lead to improvements in several areas relevant to Alzheimer’s symptoms. For instance, rescuing sleep alterations via chemogenetic manipulation of reticular nucleus activity in AD mice resulted in reduced accumulation of Aβ (Jagirdar et al., 2021), highlighting the putative importance of sleep-based neuroprotective interventions. Moreover, numerous studies showed that melatonin, a circadian rhythm–regulating hormone, plays a neuroprotective role against AD neuropathology (Hossain et al., 2019; Hossain et al., 2021; Lin et al., 2013).

Regarding mechanistic links between AD pathology and sleep, research has shown that release of Aβ is driven by neuronal hyperactivity (Bero et al., 2011; Cirrito et al., 2005; Nitsch, Farber, Growdon, & Wurtman, 1993) and concentration of Aβ in the brain parenchyma increases during wakefulness while decreasing during sleep (Kang et al., 2009; Musiek & Holtzman, 2016). In fact, not only the reduction of Aβ release, but also the clearance of it from the interstitial space via glymphatic pathway appears to be facilitated by sleep, particularly associated with elevated SWA (Iliff et al., 2012; Mendelsohn & Larrick, 2013). Altogether, this evidence suggests that improving sleep quality may thus enhance the brain’s ability to reduce Aβ load, potentially offering a therapeutic strategy for mitigating Alzheimer’s risk and/or slowing its progression.

Currently, there are no universally accepted effective treatments able to slow AD progression and relieve cognitive symptoms in patients. Sleep, contrasting to other prominent risk factors and/or outcomes of AD pathology, such as ApoE genotype, brain atrophy, or decreased cerebral blood flow, is a modifiable variable of relatively easy access. Thus, restoring sleep quality is a promising target to alleviate disease pathology and symptoms and modulating SWA during NREMS presents a valuable research opportunity with potential clinical application. However, there is a lack of interventional studies enhancing sleep during different progression stages of AD, which are crucial to determine whether sleep-based interventions may slow or stop disease progression. Here, aiming to determine the window of opportunity for sleep-based therapeutics to effectively ameliorate AD hallmarks, we investigate the effect of 2-week sodium oxybate (SO) oral administration on SWA electroencephalographic (EEG) measures, amyloid pathology, and behavioral performance in a mouse model of AD in two distinct disease stages: plaque free (early intervention) and plaque burdened (late intervention).

## Material and methods

### Animals

We used male and female Tg2576 mice (age 6 months old for early intervention, and 11 months old for late intervention) overexpressing a mutant form of amyloid precursor protein (APP), APPK670/671L, linked to early-onset familial AD, and non-transgenic WT mice from the same strain (Taconic Biosciences; Cologne, Germany). The animal room temperature was constant at 21 – 23 °C, with a 12:12 h light:dark cycle. Mice had access to food and water *ad libitum* and daily routine health checks throughout the study. The study was conducted with approval of the Cantonal Veterinary Office Zurich under license ZH210/17.

### Experimental design

All mice underwent electroencephalography/electromyography (EEG/EMG) implantation surgery two weeks prior to the start of the intervention periods (early and late intervention, **Figure 1**). Subjects were randomly allocated to the placebo or the SO treatment group, within each genotype. Baseline (BL) cognitive ability and 24-h sleep/wake behavior were assessed before the treatment period started. SO (300mg/kg, p.o.) and placebo treatments were administered twice daily (light period’s 1^st^ and 8^th^ hour) for two weeks (weekdays ON, weekends OFF treatment). Twenty-four-hour sleep/wake recordings were re-evaluated during treatment and cognitive ability was re-tested after the treatment period was concluded. Lastly, the mice were euthanized, and the brains harvested for further molecular and histological analyses.

**Figure 1.**
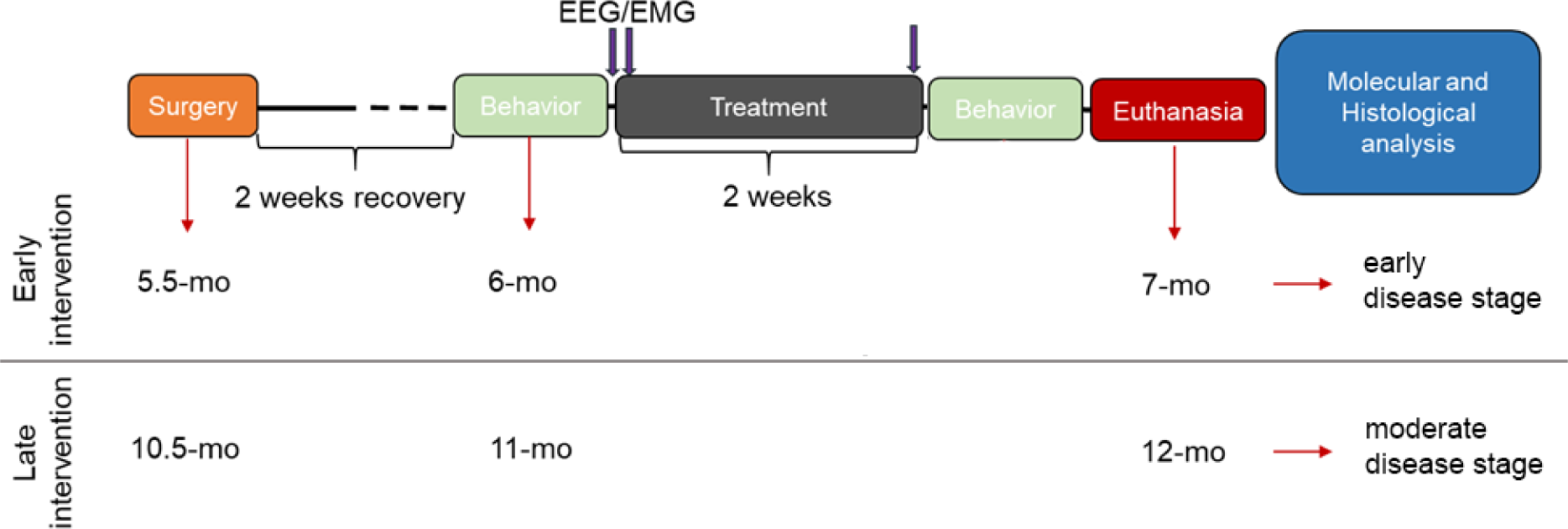
Schematic experimental design. Two weeks after EEG/EMG electrode implantation surgery (recovery period), training and posterior evaluation of baseline cognitive ability in the T-maze forced alternation test (behavior) took place, followed by baseline EEG/EMG 24-h recordings. Next day, treatments with either placebo or SO (300mg/kg, p.o.) started in combination with 24-h EEG/EMG assessment under treatment. On the last day of the two-week treatment (5 days ON, 2 days OFF), 24-h EEG/EMG assessment under treatment took place again, followed by behavioral performance re-testing and euthanasia of all experimental subjects for brain harvesting for posterior histological and molecular assessments. EEG/EMG: electroencephalography/electromyography; mo: months old.

### EEG/EMG implantation surgery

We performed all surgical procedures under deep inhalation anesthesia with isoflurane (4 - 4.5% for induction in anesthesia box, 2 - 2.5% for maintenance using a nose cone fitting), and applied lidocaine, a local anesthetic (Xylocaine, Zurich) on the surface of the head skin prior to surgery. At the end of the surgery, we administered a combination of an anti-inflammatory (5 mg/kg, s.c., Metacam®, Zurich) and a pain relief drug (0.1 mg/kg, s.c., Temgesic®, Zurich), to further prevent postoperative inflammation and pain.

The animals were implanted as described before (Buchele et al., 2016). Briefly, we placed 2 stainless steel screws (Bossard, #BN650, 1421611), one for each hemisphere, located 2 mm posterior to Bregma, 2 mm lateral from midline. We then inserted two gold wires bilaterally into the neck muscles, and applied sutures to close the skin around the implant. We connected the peridural screws and the muscle leads to a pin header (Farnell, #M80-8530445) for EEG and EMG recording, and fixed the structure with dental cement. Postsurgical analgesia was administered throughout the three days following the surgery during both light (0.1 mg/kg, s.c., Temgesic®, Zurich) and dark (1 mg/kg via drinking water, Temgesic®, Zurich) periods. We monitored wound healing, body weight and home cage activity on a daily basis during the first week after the surgery and once per week thereafter.

### Data acquisition, EEG scoring and post-processing

EEG/EMG recordings took place after a ∼48 h habituation to recording conditions in all mice. The mice were tethered to a rotating swivel and had access to every corner of the recording chamber. Signals were amplified using an N700 polysomnography (PSG) amplifier (Embla, Ontario, Canada) with a bipolar montage, digitized and collected at a sampling rate around 200 Hz and an input range of ±200 mV using a 16-bit digital-to-analog-converter. Data was collected via Somnologica Science, version 3.3.1 (Resmed, Saint-Priest, France). The scoring of the vigilance states was determined using SPINDLE (Miladinovic et al., 2019). Post-processing of EEG signals was carried out using MATLAB (ver. R2019b) as described before (Kollarik et al., 2022). Briefly, we first removed artifacts by detecting clipping events, followed by a 3-point moving average and a basic Fermi window function. Next, we resampled EEG signals at 200 Hz and filtered them between 0.5 – 30 Hz using low- and high-pass zero-phased equiripple FIR filters (Parks–McClellan algorithm; applied in both directions, filtfilt). We inspected the signal for regional artefacts not detected during automatic scoring or the above-mentioned preprocessing steps. Within scored NREMS, brief portions of the signal (<7 sample points at 200 Hz) > ± [8× interquartile range] were reconstructed by piecewise cubic spline interpolation from the neighboring points. Finally, we extracted measures of power in specific bandwidths, by performing a spectral analysis of consecutive 4-s epochs (FFT routine, Hamming window, 2-s overlap, and resolution of 0.25 Hz) and normalized the data indicating the percentage of each bin with reference to the total spectral power between 0.5 and 30 Hz.

### T-maze forced alternation test

We trained the mice in a rewarded alternation T-maze test adapted from (Deacon & Rawlins, 2006). First, we fed the mice with the reward in the home cages to habituate them to its taste and eliminate hyponeophagia. From the day prior to test until the end of the test, all animals were restricted to eat 1.5 g/mouse/day of regular chow to increase the motivation for the reward. We split the 1.5g of food into small pieces in order to avoid monopolizing of the food by the dominant mouse in the cage. On the second day, we habituated the mice to the enclosed maze by raising all doors, filling the food wells with reward, and placing at once an entire home cage group in the maze for 3 min. After the habituation day, we trained each subject in 3 trials per day for two days, with an inter-trial interval of ca. 20 min for each mouse. Each trial consisted of a two parts: the first one was a forced attempt, in which one of the arms was blocked by a door constraining the mouse to choose the open arm. The second part of the trial consisted of a choice attempt that started after the mouse consumed the reward in the open arm – or explored the arm for 2 min – and was placed back in the starting point facing away from the goal arms and forcing door was raised. Thus, in the second half of the trial, two goal arms were opened, but only the previously blocked arm had the food reward. Entering the previously not explored, now rewarded arm counted as “right” response and entering the initial arm counted as “wrong” response. On the experimental day, each mouse was subjected to 4 trials.

### Immunohistochemistry and stereology

After fresh harvesting, the brains were split into left and right hemi-brains. A randomly chosen hemi-brain was post-fixed in 4% paraformaldehyde in phosphate saline buffer (PBS), and the remaining hemi-brain was placed in an 1.5 ml Eppendorf tube and immediately snap frozen in liquid nitrogen. The post-fixed hemi-brains were dehydrated in sequential 15% and 30% sucrose in PBS solutions, then embedded in 30% sucrose moulds, and then cut into 40µm thick sagittal sections using a freezing stage-equipped microtome (Leica SM2000 R). Every 6th section was selected and mounted onto gelatinized superfrost ultra plus slides and dried at 37℃ overnight, amounting to a total of 5 sections per mouse.

#### 4g8 Staining

The mounted sections were washed in Tris-HCl buffered saline (TBS), incubated in 70% formic acid for 15 minutes, washed again in TBS, and incubated in 0.6% hydrogen peroxide (H2O2) for 30 minutes at room temperature to quench endogenous peroxidase activity. Then, sections were rinsed in TBS and blocked in M.O.M blocking reagent (Vector Laboratories, BMK-2202) diluted in 0.3% Triton-X in TBS (PH: 7.5-7.8) at room temperature for 1 h. After rinsing in TBS, the sections were first incubated for 10 min in working solution of M.O.M Diluent at room temperature and then in primary antibody (1:2000, 4g8 antibody, Covance, Cat. No. 800701, diluted in M.O.M Diluent) overnight at 4°C with gentle shaking. The following day they were incubated in M.O.M. Biotinylated Anti-Mouse IgG Reagent (Vector Laboratories) for 15 min and then in freshly prepared ABC elite solution for 1 h. After washing first in TBS and then with Tris—based buffered saline (TB) (pH = 7.6), the sections were developed in DAB (0.025%) and H2O2 (0.05%) in TBS for 25 min. Reaction was stopped by washing them three times for 10 min in TB. Subsequently, the slides were immersed in a series of ethanol solutions of increasing concentrations until 100%, then in a Xylene substitute (Roticlear®) and finally coverslipped with mounting reagent (Rotimount®).

#### Congo red Staining

Congo red staining was conducted as previously described for cerebral amyloid angiopathy (Wilcock, Gordon, & Morgan, 2006). Briefly, the mounted sagittal brain sections were first rehydrated by immersing the slides into distilled water for 30 seconds. This was followed by a two-step incubation process: first, the sections were placed in an alkaline saturated NaCl for 20 minutes, then incubated in Congo red solution for 40 minutes. The slides were rinsed briefly in 95 and 100% ethanol solution and subsequently treated with three series of Xylene, each for 5 minutes. Finally, the slides were cover-slipped with mounting reagent.

#### Stereology

Five sagittal sections per mouse were used for stereological estimations. The area fraction fractionator probe (Stereoinvestigator™, MBF Bioscience) was used to estimate the areas covered by plaques in the regions of interest (ROI). The plaques were visualized with a Zeiss Imager M2 using a 20x objective, the size of counting frame was 150 x 150 μm for hippocampus and 200 x 200 μm for cortex. Random sampling grid size was identical to the counting frame size in order to scan the entire ROI.

### Brain homogenization and ELISA

We mechanically homogenized the frozen hemi-brains and sequentially extracted soluble and insoluble proteins with diethylamine (DEA) and formic acid (FA), respectively (Casali & Landreth, 2016). Aβ content in soluble and insoluble brain extracts was assessed by sandwich ELISA as indicated by the kit’s manufacturer (Aβ-40: KMB3481, Aβ-42: KMB3441, Thermo Fischer). To serve as controls the placebo-treated Tg2576 brains were blindly processed in identical fashion and in parallel to SO-treated Tg2576 brains throughout the entire process. Soluble and insoluble Aβ-40 and Aβ-42 were normalized to brain tissue mass and expressed in pg/mg.

### Statistical analyses

Data analyses were carried out using IBM^®^ SPSS^®^ Statistics 25 software and the data were visualized by GraphPad Prism 9 (GraphPad Software, Inc., San Diego, CA). Outliers were detected with boxplots, normality was assessed using Skewness and Kurtosis for each variable, and homogeneity of variances was assessed with Levene’s test. Delta activity gain was analyzed with multiple t-tests for each hour and *p-*values were corrected with false discovery method in GraphPad Prism 9. NREMS proportion was analyzed with two-way ANOVA (genotype*treatment). Analyses of the main effect of genotype and treatment were separately run and followed by pairwise comparisons. All pairwise comparisons were reported with 95% confidence intervals (CI) and *p*-values. Cognitive performance of the mice was assessed in two steps. First, we conducted independent t-tests to investigate their baseline differences between genotypes in each age group, separately. Second, we ran paired t-test to evaluate the change in cognitive performance from baseline to after treatment in Tg2576 mice. Biserial correlation analysis was used to assess the relationship between delta activity gain and cognitive performance in Tg2576 mice after cognitive performance was assigned to two categories: fail (≤ 50) or success (> 50). Briefly, we first computed point-biserial correlation in SPSS and then transformed the point-biserial correlation coefficient (r_pb_) into biserial correlation coefficient (r_b_) as it has been described before (Field, 2012). The difference in levels of soluble and insoluble Aβ-40 and Aβ-42 between placebo and SO treated Tg2576 mice was assessed with independent t-test. The plaque burden in the hippocampus and cortex was calculated by dividing the total area covered with plaques by the total area of ROI. Then, the data were analyzed with a two-way ANOVA (genotype*treatment), followed by analyses of main effects of genotype and treatment, separately. Subsequently, pairwise analyses were run to assess whether there was a difference in plaque burden between placebo- and SO-treated mice within the same genotype, and whether there is a difference between Tg2576 and WT mice within the same treatment. Pearson’s correlation analysis was run to evaluate the relationship between delta power gain and plaque burden in the hippocampus. Quantitative estimates of amyloid plaques burden in WT and Tg2576 young and aged mice identified through Congo red staining were analyzed using independent samples t-test.

## Results

### SO administration elicited delta activity gain with unaltered time spent in NREMS

Decreased delta power during sleep is associated with AD (Lee, Gerashchenko, Timofeev, Bacskai, & Kastanenka, 2020) and may be one of the factors that worsen disease symptoms (Bianchetti et al., 1995; Wennberg et al., 2017). Considering that Tg2576 mice present with reduced delta activity in NREMS (Kollarik et al., 2022), we first aimed to rescue the impaired delta activity by SO pharmacotherapy. We administered placebo or SO (300 mg/kg, p.o.) in the 1^st^ and 8^th^ hour of the light period for all groups. As first step, we evaluated whether such pharmacological regime exerted an increase in the time spent in NREMS compared to baseline. Our results demonstrate that administration of SO (300 mg/kg, p.o.) did not significantly alter the time spent in NREMS in any of the groups in comparison to their own baseline (**Figure 2A, B**). Delta activity gain, i.e. increased SWA, from baseline was especially observed after the administration in the 8^th^ hour in all groups treated with SO when compared with the placebo-treated groups (**Figure 2C-F**). A difference in delta activity gain between SO and placebo after the 1^st^ hour administration was apparent only in Tg2576 mice in the late intervention cohort (**Figure 2F**).

**Figure 2.**
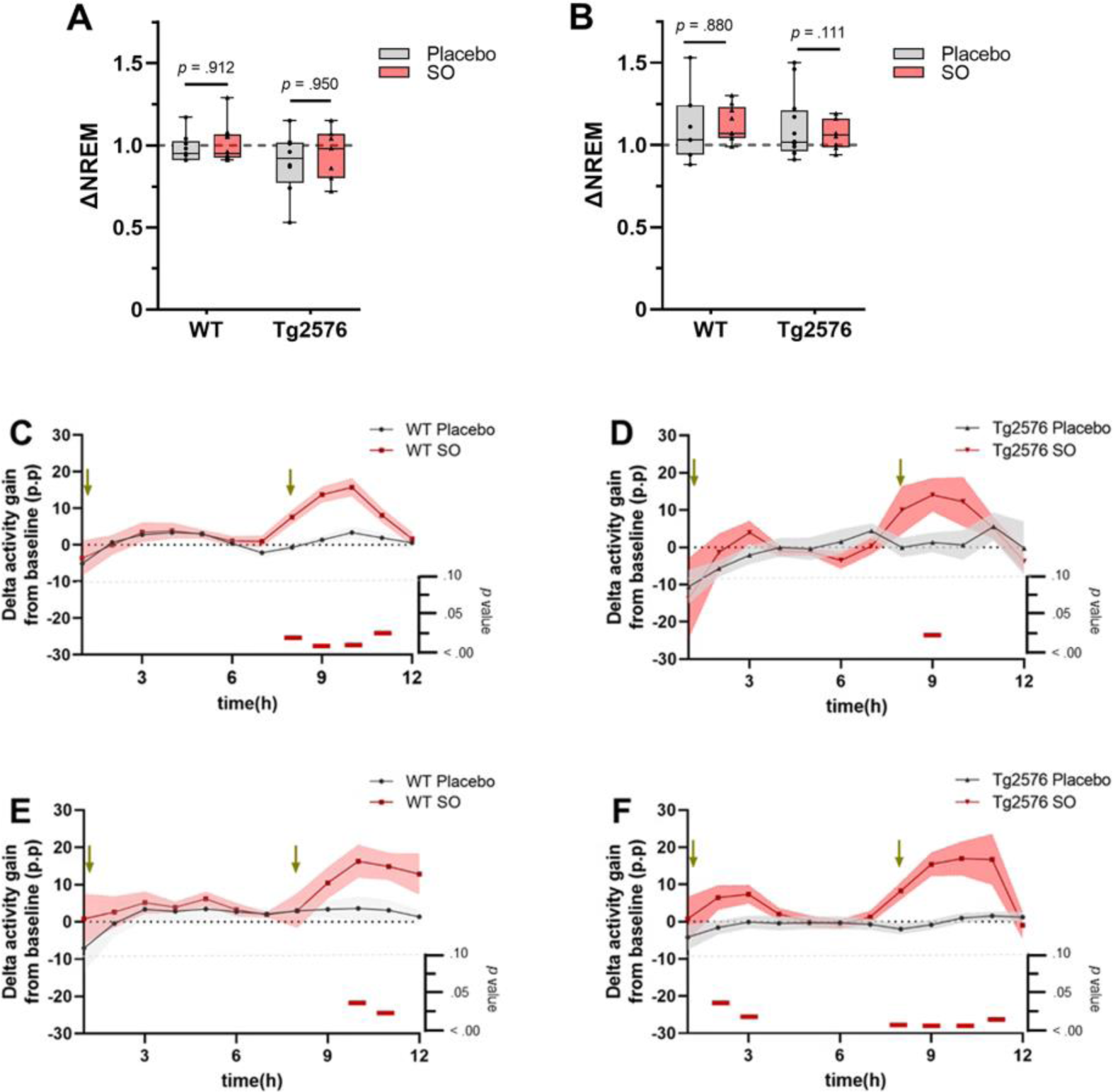
NREMS proportion and delta activity gain upon placebo or SO treatment in early and late disease stage WT and Tg2576. **A)** Placebo and SO treatments did not change the time spent in NREMS compared to baseline (ΔNREM) neither in either early, or **B)** late intervention groups. Data is expressed as mean ± standard error (SE-), two-way ANOVA (genotype*treatment). **C)** Strong evidence of delta activity gain from baseline was present after SO administration (red) in the 8th hour in both early intervention WT, and **D)** Tg2576 mice, as well as **E)** late intervention WT, and **F)** Tg2576 mice compared to age-matched placebo (gray) treated groups. Green arrows indicate times of treatment administration. Data is expressed as mean ± SE-, multiple t-test. *P* values are indicated in the bottom of each figure, with horizontal red bars highlighting significant time bins. Δ: change; WT: wild-type; Tg2576: AD mice; SO: sodium oxybate; p.p.: percentage points.

### SO treatment improves cognitive performance in mice at early stage of AD

Studies showed that memory loss starts around 6 months of age in Tg2576 mice, which coincides with the appearance of detergent-insoluble Aβ aggregates (Westerman et al., 2002). Therefore, we first assessed cognitive performance of Tg2576 mice and WT controls in the T-maze forced alternation test in both age groups, separately, to determine the existence of treatment-naïve (baseline, BL) differences in memory (**Figure 3A-B**). We observed that Tg2576 mice in both age groups (early intervention: n = 17, M = 42.65, SD = 14.99; late intervention: n = 20, M = 53.75, SD = 29.43) performed poorly (early intervention: *t*(26.21) = 4.884, *p* < .001, Cohen’s *d* = 1.64; late intervention: *t*(36) = 2.753, *p* = .009, Cohen’s *d* = 0.91) compared to age-matched WT controls (early intervention: n = 18, M = 79.63, SD = 28.18; late intervention: n = 18, M = 75.93, SD = 18.28) in the T-maze test.

**Figure 3.**
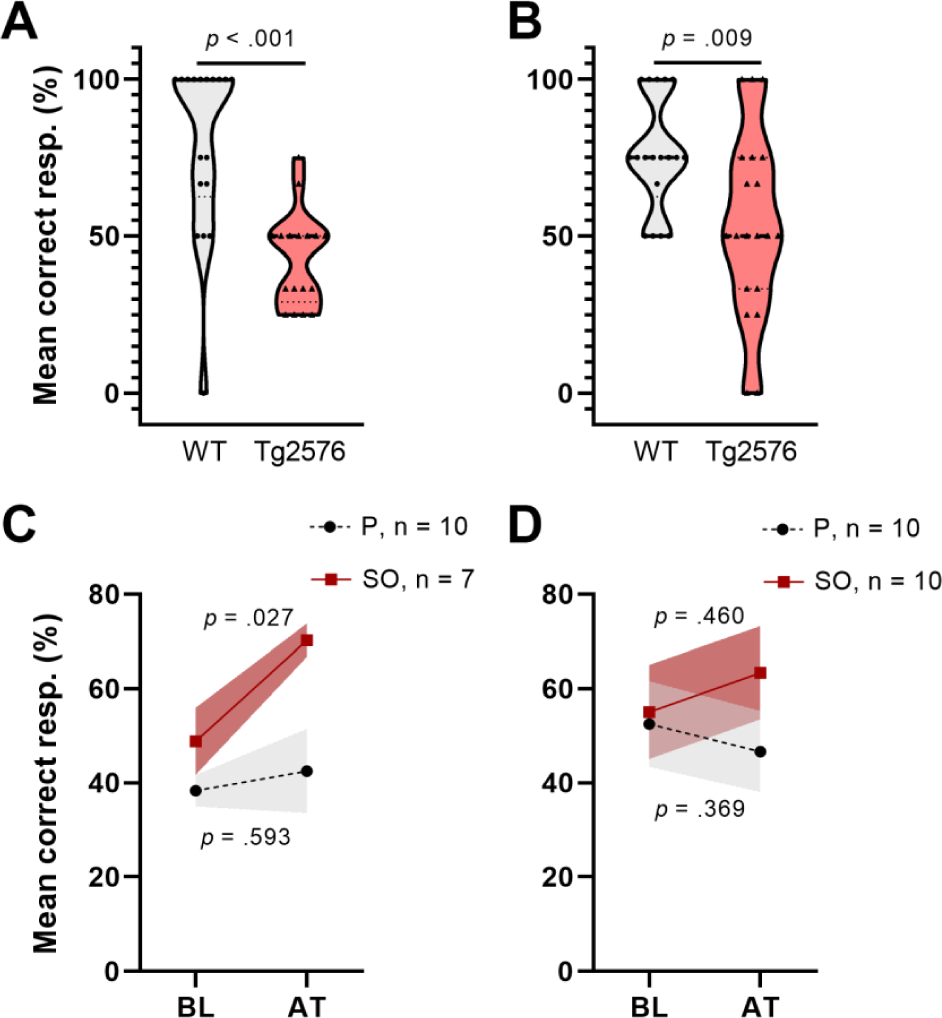
T-maze forced alternation test performance of early and moderate disease stage Tg2576 mice at baseline and upon treatment. **A)** Tg2576 mice performed poorly in both early and **B)** moderate disease stage in comparison to age-matched WT mice (Independent t-test; early: p<0.001; late: p=0.009). **C)** Early disease stage Tg2576 mice significantly improved performance values after two weeks of treatment with SO (paired t-test, *p* = .027), whereas placebo treated mutants draw no benefit from the intervention (paired t-test, *p* = .593). **D)** Moderate disease stage Tg2576 mice do not present significantly improved performance scores upon treatment (paired t-test, *p* =.46), same as their placebo-treated counterparts (paired t-test, *p* =.369). Data is expressed as mean ± SE-. resp.: responses; WT: wild-type; Tg2576: AD mice; P: placebo; SO: sodium oxybate.

On the other hand, increases in high amplitude SWA during NREMS may be associated with improved cognition (Wilckens, Hall, Nebes, Monk, & Buysse, 2016). Thus, we analyzed Tg2576 mice cognitive performance in both age groups after treatment (AT) with placebo or SO (**Figure 3C-D**). Our analyses revealed that cognitive performance in the early intervention SO group improved AT (**Figure 3C**) (BL: M = 48.81, SD = 18.90; AT: M = 70.24, SD = 9.45; *t*(6) = 3.286, *p* = .017, Cohen’s *d* = 1.43), while treatments with placebo did not affect this outcome (BL: M = 38.33, SD = 10.54; AT: M = 42.50, SD = 28.18; *t*(9) = 0.460, *p* = .657, Cohen’s *d* = 0.20). There were no significant effects of either placebo (BL: M = 52.50, SD = 28.61; AT: M = 46.67, SD = 27.27; *t*(9) = −0.771, *p* = .460, Cohen’s *d* = 0.16) or SO (BL: M = 55.00, SD = 31.72; AT: M = 63.33, SD = 31.23; *t*(9) = 0.905, *p* = .389, Cohen’s *d* = 0.26) over Tg2576 mice cognitive ability in the late intervention group (**Figure 3D**).

### Cognitive performance is positively correlated with SO-triggered delta activity gain

After determining increased delta activity and improved cognitive performance in Tg2576 mice after 2 weeks of oral SO administration, we examined whether there is an association between both measures. To this end, we first assigned the cognitive performance scores as fails (≤ 50%) or successes (> 50%), and then ran a biserial correlation between treatment-elicited delta activity gain after second administration (delta gain in the 9^th^ + 10^th^ hours) in both treatments groups, i.e. placebo and SO, and cognitive performance in Tg2576 mice of both age groups, separately. Our analyses showed a trend towards positive correlation between delta activity gain and cognitive performance in the early intervention group (n = 8, r_pb_(6) = 0.653, r_b_ = 0.819, *p* = .079, **Figure 4A**), while there was no apparent correlation in the late intervention group (n = 9, r_pb_(7) = 0.563, r_b_ = 0.786, *p* = .114, **Figure 4B**).

**Figure 4.**
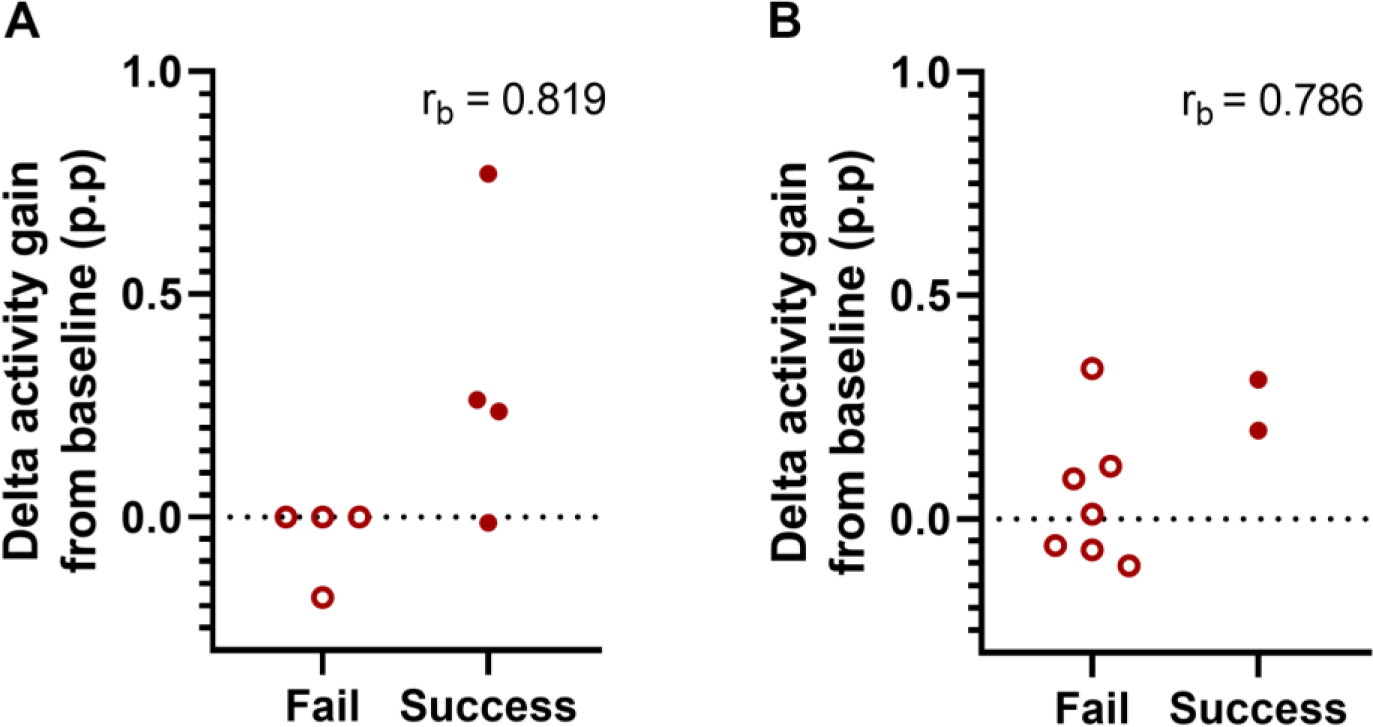
Association between delta activity gain and cognitive performance in early and moderate disease stage Tg2576 mice. **A)** Biserial correlation analyses revealed a positive large correlation coefficient between delta activity gain and the cognitive performance in both early and **B)** late intervention cohorts. No significant results were identified due to small sample size. 0.1 < | r_b_ | < .3 = small correlation, 0.3 < | r_b_ | < .5 = medium/moderate correlation, | r_b_ | > .5 = large/strong correlation. p.p.: percentage points.

### Histopathological evaluation of the baseline conditions in early and late intervention groups

Despite the progressiveness of AD pathology in Tg2576 mice has been widely reported before (Kawarabayashi et al., 2001), differences in the time onset suggested across individual colonies may lead to uncertain baseline conditions for longitudinal histopathological assessments. To determine the exact pathological stage in terms of plaque formation in the 6 and 11 months old Tg2576 mice used in our study, as basis for additional assessments of the effect of our treatment onto pathology, we performed Congo red staining of sagittal brains sections from non-treated WT and Tg2576 mice across both age groups and quantitatively assessed plaque burden in the hippocampus (**Figure 5**). Representative photomicrographs from 11-month old Tg2576 mice (**Figure 5A-E**) demonstrated a high level of plaque deposition in hippocampal areas. Stereological analyses confirmed a very low and indifferent plaque burden in WT and Tg2576 mice at early disease stage (t(6) = 0.169, p = .871, Cohen’s d = 0.12, **Figure 5F**), while evidencing a higher plaque burden in Tg2576 at moderately advanced disease stage (t(7) = −3.850, p = .006, Cohen’s d = 2.50, **Figure 5G**) compared to age-matched WT littermates.

**Figure 5.**
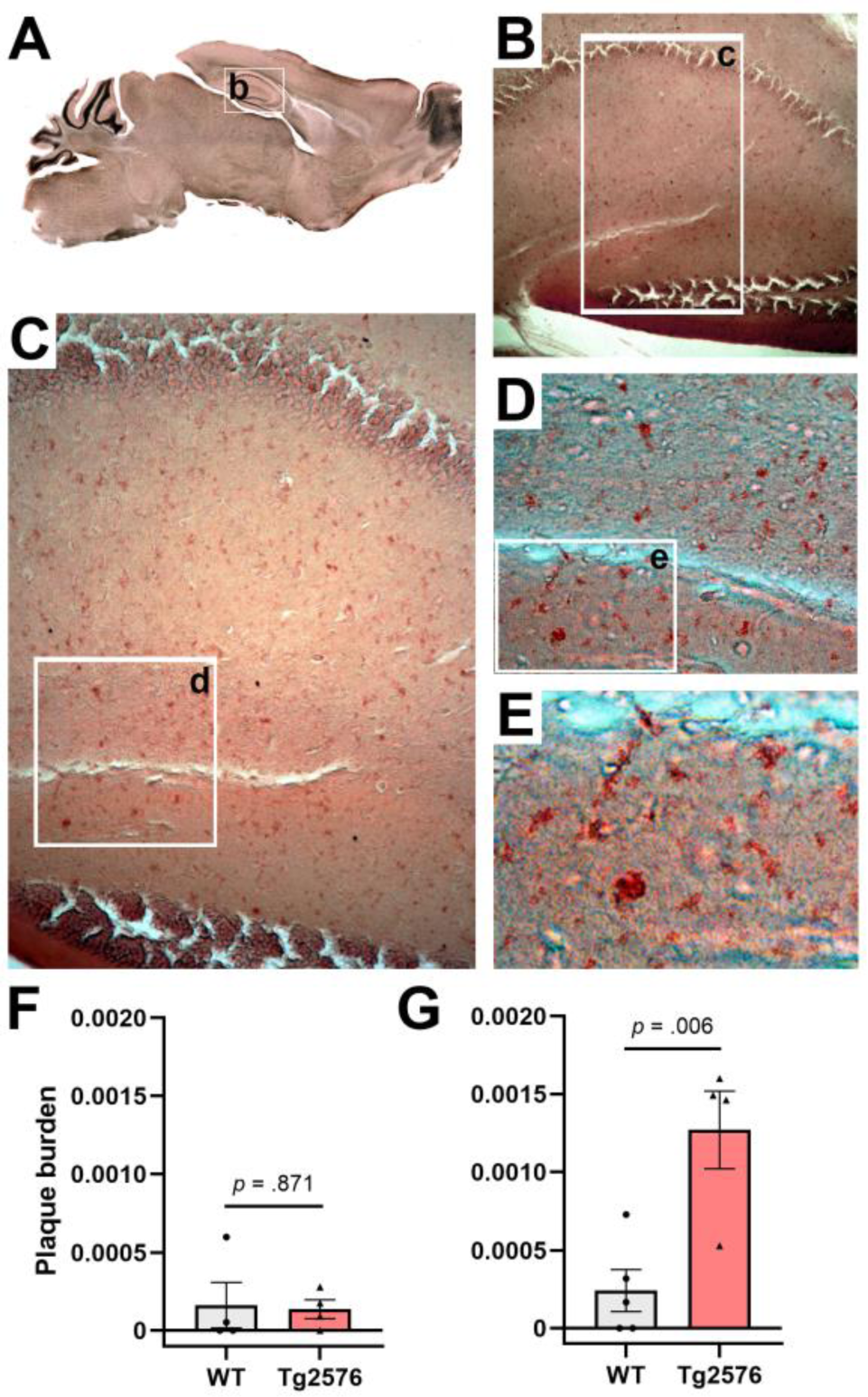
Congo red staining of hippocampal amyloid plaques in drug-naïve early and moderate disease stage WT and Tg2576 mice. **A-E)** Representative photomicrographs illustrating Congo red positive deposits in hippocampal tissue of moderate disease stage Tg2576 mice. **F)** Early disease stage Tg2576 mice and age-matched WT controls present low and indistinct levels of plaque burden in the hippocampus (independent t-test; *p* =.871), whereas **G)** moderate disease stage Tg2576 mice have a significantly higher plaque burden score than their age-matched WT controls (independent t-test; p=0.006). WT: wild-type; Tg2576: AD mice.

### SO treatment results in reduced levels of toxic insoluble Aβ42/40 ratio and diminished plaque burden in early- and late-stage Tg2576 mice

Based on the quantification results of the Congo red staining showing now discernible plaque load in the Tg2576 mice at 6 months of age, agreeing with previous literature (Kawarabayashi et al., 2001), we proceeded to evaluate the effect of SO and placebo treatments onto amyloidosis in this premorbid (plaque free) stage via determinations of soluble and insoluble Aβ-40 and Aβ-42 levels in brain tissue from Tg2576 mice via ELISA. We observed that soluble Aβ-40 (*t*(8) = 0.638, *p* = .791, Cohen’s *d* = 0.18) and Aβ-42 (*t*(8) = 0.280, *p* = .864, Cohen’s *d* = 0.11, **Figure 6A, B**) did not differ between treatments. Insoluble Aβ-40 level, on the other hand, was higher (*t*(8) = 2.846, *p* = .022, Cohen’s *d* = 1.80) in the brains of SO-treated Tg2576 mice (n = 5, M = 4.982, SD = 0.379) in comparison to what was observed in the brains of placebo-treated Tg2576 mice (n = 5, M = 4.459, SD = 0.158, **Figure 6D**). Insoluble Aβ-42 levels did not differ (*t*(8) = 0.597, *p* = .567, Cohen’s *d* = 0.38, **Figure 6E**) between treatments. Aβ-42/Aβ-40 ratio is considered a more accurate way of reflecting the progress of AD (Wiltfang et al., 2007) than Aβ-40 and Aβ-42 expressed separately. Therefore, we followed our analyses with the assessment of treatment-dependent changes in Aβ-42/Aβ-40 ratio. We did not detect a difference in the soluble Aβ-42/Aβ-40 ratio between treatments (*t*(8) = 0.695, *p* = .723, Cohen’s *d* = 0.23, **Figure 6C**), while our results indicated a trend with a large effect size towards a lower insoluble Aβ-42/Aβ-40 ratio in the brains of SO-treated Tg2576 mice (*t*(8) = 2.009, *p* = .079, Cohen’s *d* = 1.27, **Figure 6F**) compared to placebo-treated controls.

**Figure 6.**
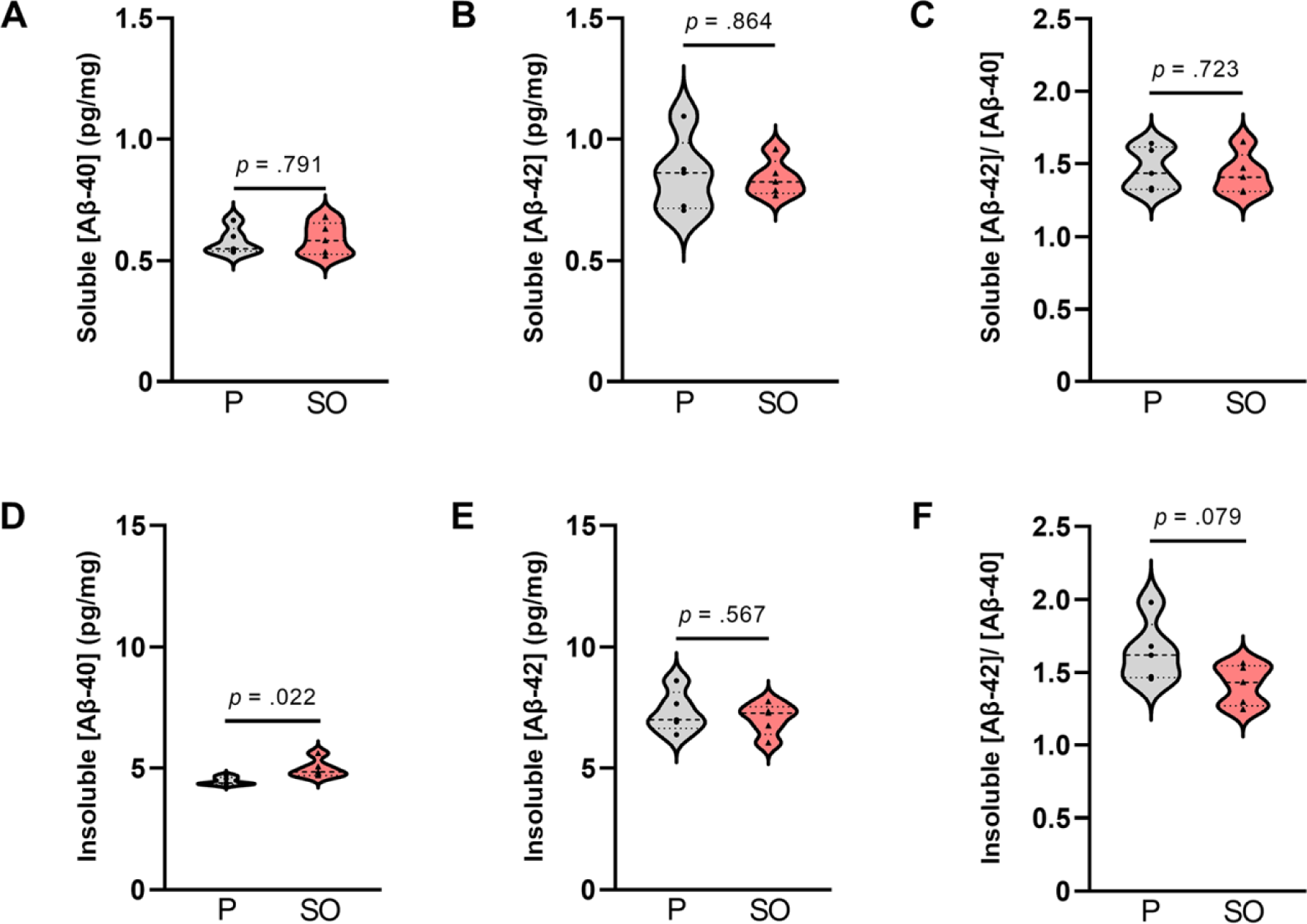
Soluble and insoluble Aβ-40 and Aβ-42 brain concentration upon treatments in early disease stage Tg2576 mice. **A)** No differences in soluble [Aβ-40], **B)** [Aβ-42], and **C)** [Aβ-42/ Aβ-40] ratio between placebo- and SO-treated early disease stage Tg2576 mice’ brains (independent t-test; Aβ-40: *p* = .791, Aβ-42: *p* =.864, Aβ-42/ Aβ-40: *p* =.723). **D)** Insoluble [Aβ-40] was higher in the brains of SO-treated early disease stage Tg2576 mice in comparison to placebo-treated mutants (independent t-test; *p* = .022), whereas **E)** insoluble [Aβ-42] did not differ between treatments (independent t-test; *p* = .567). **F)** The ratio of insoluble [Aβ-42/ Aβ-40] in SO-treated early disease stage Tg2576 mice’ brains was slightly lower than that of placebo, with a trend statistical significance (independent t-test; p=0.079). Data is expressed as mean ± SE-. [Aβ-40]: amyloid beta 40 concentration; [Aβ-42]: amyloid beta 42 concentration; P: placebo; SO, sodium oxybate.

Tg2576 mice have been reported to develop Aβ plaques as they age (Benzing et al., 1999; Hsiao et al., 1996; Ishii, Wang, Racchumi, Dyke, & Iadecola, 2014; Kawarabayashi et al., 2001), as confirmed by the significant difference in the load of Congo red-stained plaques between WT and Tg2576 mice at 11 months of age. In fact, hippocampus and cortex are two primary regions vulnerable to plaque accumulation in AD (Bero et al., 2011). Therefore, we investigated plaque burden in both these key areas in brains from the late intervention cohort treated with placebo or SO. We included WT mice in our analyses as negative controls. We found strong evidence of reduced plaque burden in hippocampal (**Figure 7A-C**) and cortical (**Figure 7D-F**) brain areas of Tg2576 mice after 2 weeks of treatment with SO. The interaction effect between genotype and treatment on plaque burden in the hippocampus (**Figure 7C**) was a trend with a large effect size (*F*(1, 28) = 3.807, *p* = .061, η_p_ = .120). The main effect of genotype on plaque burden was significant (*F*(1, 28) = 10.820, *p* = .003, η_p_ = .279), while the main effect of treatment was a trend (*F*(1, 28) = 3.634, *p* = .067, η_p_ = .115). Following pairwise comparisons within the genotype, we observed a significant decrease in plaque burden in SO-treated Tg2576 mice, while there was no change in the placebo treatment group. There was no effect of treatment on the naturally low plaque burden in the hippocampus of WT mice. Notably, after 2 weeks of SO treatment, plaque burden in the hippocampus of Tg2576 mice was at a similar level than the plaque burden in WT mice. In the cortex, plaque burden analyses revealed similar results to those in the hippocampus (**Figure 7F**). The interaction effect between genotype and treatment on plaque burden in the cortex was a trend with large effect size (*F*(1, 28) = 4.054, *p* = .054, η_p_ = .126). There was a significant main effect of genotype (*F*(1, 28) = 13.965, *p* = .001, η_p_ = .333) but not of treatment (*F*(1, 28) = 2.951, *p* = .097, η_p_ = .095) on plaque burden. We followed the analyses with pairwise comparisons and observed a greater plaque burden in placebo-treated Tg2576 mice in comparison to SO-treated mutants. Within the placebo-treated mice, Tg2576 mice showed higher plaque burden in the cortex than WT, while SO treated mice did not differ between genotypes.

**Figure 7.**
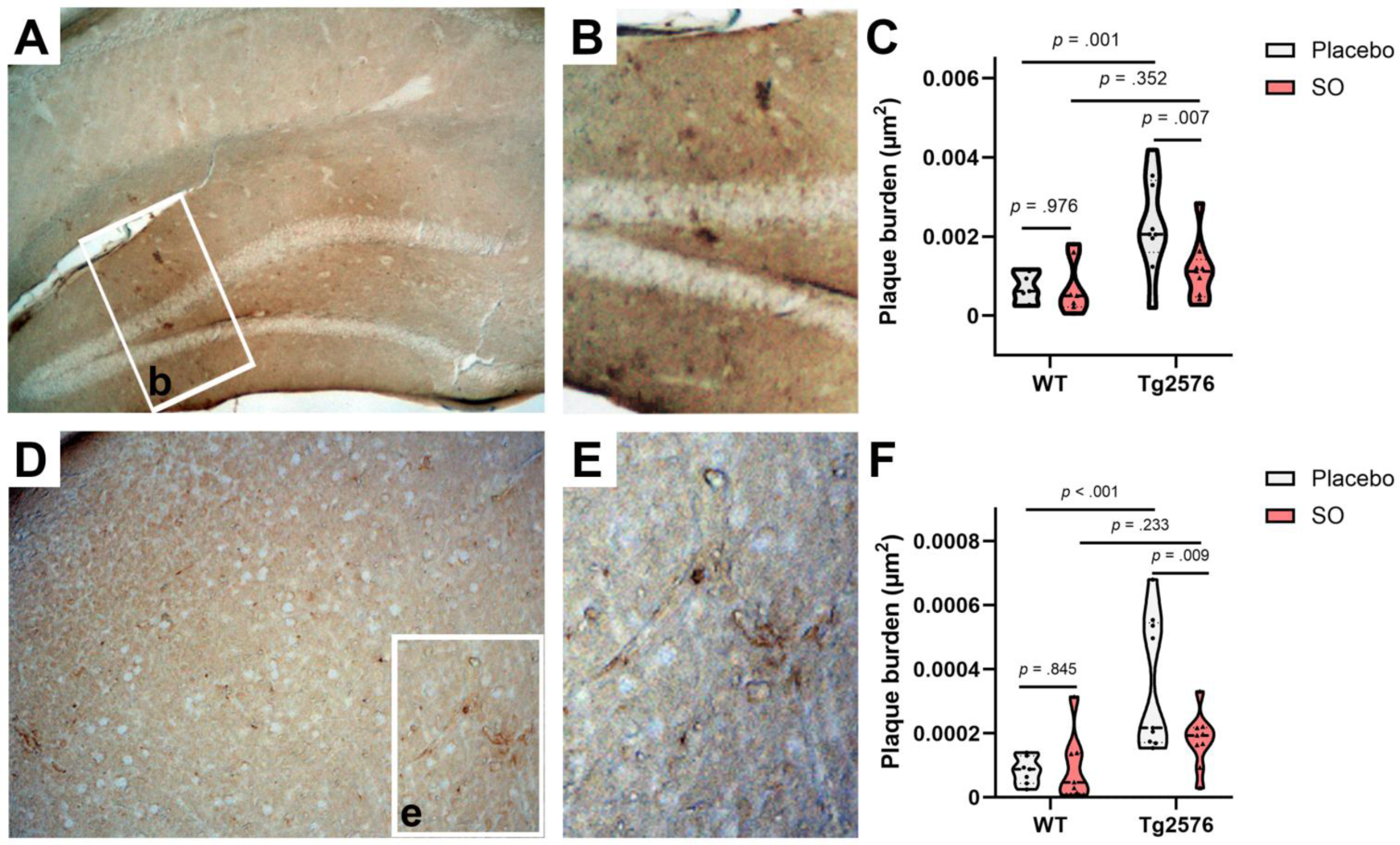
Hippocampal and cortical amyloid plaque burden upon treatments in moderate disease stage Tg2576 mice. **A, B)** Representative photomicrographs of 4g8-positive deposits in the hippocampus of moderate disease stage Tg2576 mice’s brains. **C)** Stereological estimates of plaque burden (µm^2^) in placebo-treated WT (n=7), SO-treated WT (n=7), placebo-treated Tg2576 (n=9), and SO-treated Tg2576 (n=9) mice’ brains stained against Aβ (17-24) with 4g8 antibody. A significant decrease of plaque burden was observed in hippocampus of Tg2576 brains after two-week treatment with SO compared to placebo-treated mutants (two-way ANOVA; pairwise comparisons, *p* = .007). **D, E)** Representative photomicrographs of 4g8-positive deposits in the cortex of moderate disease stage Tg2576 mice’s brains. **F)** Stereological estimates of plaque burden (µm^2^) in placebo-treated WT (n=7), SO-treated WT (n=7), placebo-treated Tg2576 (n=9), and SO-treated Tg2576 (n=9) mice’s brains stained against Aβ (17-24) with 4g8 antibody. A significant decrease of plaque burden was observed in cortical regions of Tg2576 brains after two-week treatment with SO compared to placebo-treated mutants (two-way ANOVA; pairwise comparison, *p* = .009). All data are expressed as mean ± SE-. WT: wild-type; Tg2576: AD mice; SO, sodium oxybate.

### Reduction in plaque burden is associated with SO-triggered delta activity gain

Our findings of increased delta activity gain and decreased plaque burden in Tg2576 mice after treating with SO led us to investigate the relationship between both parameters. Therefore, we ran a Pearson’s correlation between the plaque burden in the hippocampus and cortex (**Figure 8**), and delta activity gain after the second administration of the treatment (delta activity gain in the 9^th^ + 10^th^ hours). Our findings demonstrated that mice that had higher delta activity gain showed less plaque burden in the hippocampus (r = −0.882, *p* = .020; **Figure 8A**), while no association was found for cortical plaque burden and delta activity gain (r = −0.065, *p* = .902; **Figure 8B**).

**Figure 8.**
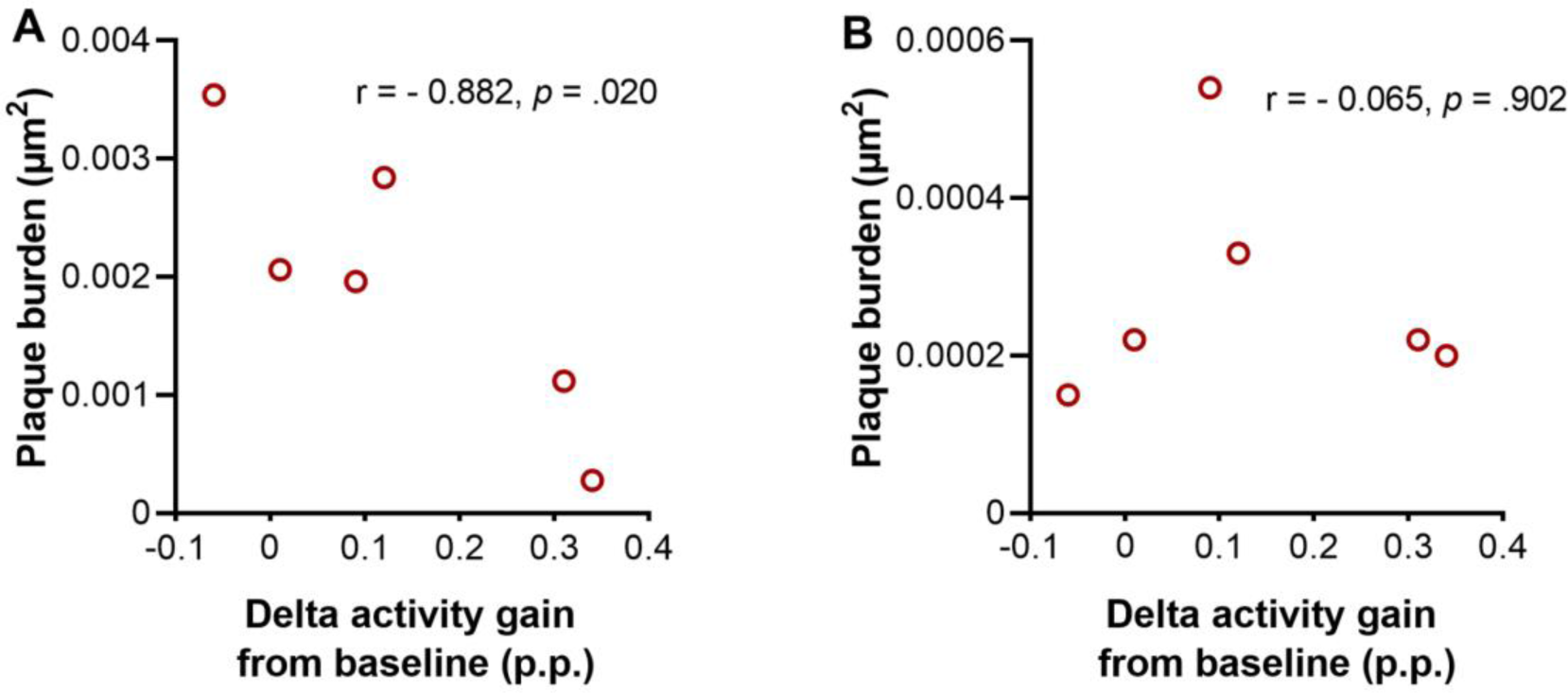
Association between delta activity gain and plaque burden in moderate disease stage Tg2576 mice. **A)** Strong negative association between plaque burden in hippocampus and delta activity gain in moderate disease stage Tg2576 mice (Pearson’s correlation coefficient r= −0.882, *p* =.020). **B)** No association between plaque burden in cortex and delta activity gain was observed in the same mice (Pearson’s correlation coefficient r= −0.065, *p* =.902). 0.1 < | r | < .3 = small correlation, 0.3 < | r | < .5 = medium/moderate correlation, | r | > .5 = large/strong correlation. p.p.: percentage points.

## Discussion

Reduced sleep quality and duration are commonly observed phenomena associated with aging. Apart from being highly prevalent in the elderly population (Mander, Winer, & Walker, 2017), sleep disturbances are often more severe and exacerbate disease symptoms in AD and other neurodegenerative diseases. Cumulating evidence supports a reciprocal relationship between sleep and AD, meaning that disturbed sleep is not only an outcome of AD but can also affect cognitive function and disease pathology in patients (Lucey et al., 2021; Wang & Holtzman, 2020). Drawing on this relationship, treatments targeting the restoration of sleep could be a promising approach to alleviating disease symptoms.

It was previously shown that enhancing sleep with a dual orexin antagonist decreased Aβ aggregation in a mouse model of AD (Kang et al., 2009). Moreover, another sleep promoting neuromodulator, melatonin, inhibited the generation and formation of amyloid fibrils *in vitro* (Lin et al., 2013) and diminished tau hyperphosphorylation (Lin et al., 2013; Ling et al., 2009). In a previous study, we demonstrated that SWA enhancement via SO up-regulates multiple proteostatic pathways with capacity to regulate intra- and extracellular noxious protein levels in Parkinson’s disease mice (Morawska et al., 2021). Moreover, optogenetic targeting of GABAergic interneurons in a mouse model of AD not only rescued sleep disruptions and sleep fragmentation by improving NREMS, delta power and SWA, but also increased microglial clearance ability resulting in phagocytic uptake of Aβ (Zhao et al., 2023). In the present study, as SWA modulator we used SO, which is a GABA_B_/GHB receptor agonist known to generate deep sleep-like increases in delta power to investigate the restorative effect of SO-induced “deep sleep” on cognition and amyloidosis in a well-established mouse model of AD, the Tg2576 line.

AD pathogenesis in humans begins 10-20 years prior to the onset of clinical symptoms (Sperling et al., 2011). Thus, it is important to establish potential disease modifying therapies targeting the preclinical phase for the prevention of AD. It has been predicted that as small as a 1 year delay of AD symptoms onset can result in 11.8 million fewer cases worldwide, massively reducing the global burden of the disease (Brookmeyer, Johnson, Ziegler-Graham, & Arrighi, 2007). Moreover, the unsuccessful clinical translation of positive results obtained from targeting the late phases of AD in animal models (Knopman, Jones, & Greicius, 2021; Tampi, Forester, & Agronin, 2021) calls for novel therapeutic strategies focusing on prevention and/or delaying the onset of AD before the full spectrum of clinical hallmarks is present. In line with this, we designed our study aiming to find an effective time course to start sleep-based treatments to potentially alleviate pathological features and cognitive symptoms of AD. Thus, we selected two relatively early age and treatment time-points, in which mice are either in the pre-morbid stage (amyloid plaque-free) or in moderate stage (amyloid plaque-burdened), which we successfully confirmed via quantitative analysis of Congo red staining.

Our findings indicate that oral SO administration with a dose of 300mg/kg increases delta activity without changing the 24-h sleep amount in both Tg2576 and WT mice, and provide evidence of reduced plaque deposition in both hippocampus and cortex in moderate disease stage Tg2576 mice. Consistent with our observations, Klein et al. showed decreased levels of Aβ due to enhanced activity of nephrilysin after oral treatments with SO (Klein et al., 2015). Therefore, our work further contributes to existing knowledge of a positive effect of “deep sleep” promotion on Aβ plaque clearance (Kang et al., 2009; Klein et al., 2015). Additionally, our correlation analysis revealed that the mice with higher delta activity gain after SO administration, also presented lower plaque burden in the hippocampus, suggesting a link between sleep depth and Aβ clearance. However, this correlation was not observed in the cortex. The most probable reason for the absence of an association between delta gain and plaque burden in the cortex is the high variability of plaque burden measures in this region, which may result from our quantification method. Future studies should consider sub-dividing the measurements into distinct cortical regions (e.g. prefrontal cortex, parietal cortex, piriform cortex) to prevent the spreading out the data.

Given the absence of stainable/visible plaques in Tg2576 mice at the age of 6-7 months (even incipient plaques are seen only at 11-12 months), we assessed amyloidosis via ELISA by investigating the levels of soluble and insoluble Aβ-40 and Aβ-42, and their respective ratios. Surprisingly, we have not found any difference in soluble Aβ between the placebo and SO treated AD mice. A possible explanation for this finding might be the 3-day OFF treatment period granted to all mice before euthanasia, likely allowing enough drug washout and consequent re-establishment of reduced levels of SWA. Reinstated low SWA could in turn lead to neuronal activity-dependent re-normalization of pathological soluble Aβ levels. This important finding suggest that a continuous treatment might be necessary to avoid re-accumulation of toxic amyloids in critical brain regions. Insoluble Aβ levels, on the other hand, revealed a significantly higher Aβ-40 level in SO treated Tg2576 mice compared to placebo-treated ones, while insoluble Aβ-42 levels did not differ between treatments. Relatively higher levels of shorter, less fibrillogenic Aβ-40 in SO-treated mice emerges as an intriguing factor with potential neuroprotective implications, also previously discussed by others (Kim et al., 2007; Zoltowska, Maesako, & Berezovska, 2016). The slight reduction with a large effect size found in Aβ-42/Aβ-40 ratio after two weeks of treatment with SO in early disease stage mice may additionally suggest prevention neurotoxicity (Zoltowska et al., 2016).

Our assessments of cognitive ability by T-maze forced alternation test revealed working memory deficits in the transgenic mice of both age groups. After two weeks of treatment with SO, cognitive performance improved in plaque-free stage Tg2576 mice, but no symptom alleviation was observed in moderate disease stage mice. Westerman et al. found that Tg2576 mice exhibited behavioral changes consistent with the acceleration of insoluble Aβ at 5–6 months of age. However, there was no association between insoluble Aβ and behavioral changes across the broader age range of 5 to 22 months.(Westerman et al., 2002). Despite clearance of Aβ plaques through SO-boosted delta power, the lack of improvement in memory performance in Tg2576 mice at moderate disease stage raises the question of whether Aβ directly influences cognition or if it rather has an indirect role in damaging neurons and in return impacting cognition by inducing neurotoxicity or inflammation. Thus, the accumulation of amyloid plaques may lead to a point where it is too late to reverse cognitive impairment, as the damage to neurons, synaptic connections, and brain networks becomes extensive and irreversible.. Whether a longer intervention period (16-week SO administration regime started but disabled half way through due to COVID19 pandemic) would result in improved cognition in moderate disease stage remains to be determined.

Correlation analyses between delta activity gain and cognitive performance in Tg2576 mice, although observed in the expected direction with strong effect sizes, were not statistically significant for either age group. Therefore, future studies shall consider larger sample sizes to strengthen particularly the behavioral analysis, as well as >1 day EEG/EMG recording for a better interpretation of these results. Others have shown before that oral administration of SO via drinking water decreased levels of brain Aβ and reduced cognitive deficits in APPSWE mice (Klein et al., 2015). A limitation of that study, however, was the difficulty to control the dose of SO consumed by the animals, hindering the possibility to draw links between specific sleep alterations elicited by the treatment and the observed effects. In fact, the study did not include objective sleep measures and, therefore, a direct link between SO administration and its major effector (i.e. enhanced SWA) were not drawn. Thus, to the best of our knowledge, ours is the first study reporting an effect of increased SWA on both amyloid pathology and cognitive symptoms in a mouse model of AD, specifically in a disease-stage dependent manner.

Several limitations our study, however, need to be taken into consideration. First, the absence of a model of tauopathy, another pathological hallmark of AD which is strongly correlated with the disease symptoms, impairs our ability to generalize our findings to AD. Evaluating the efficacy of SO treatment onto an appropriate model of tau pathology would be important for future studies. Second, we did not evaluate whether a longer period of treatment would be effective on cognitive performance of Tg2576 in the moderate disease stage group. Knowing whether longer duration of sleep treatment interventions may additionally help arrest symptoms at an advanced disease stage would be of great importance to develop potential treatments for advanced AD patients. Finally, pharmacological sleep modulators could potentially alter fine-tuned neurochemical brain systems directly (i.e., bypass sleep, their main target) and have unspecific secondary effects confounding the interpretation of results. Therefore, future studies should aim at nonpharmacological sleep modifying methods, such as auditory stimulation of sleep slow oscillations (Dias, Baumann, & Noain, 2024; I. Dias et al., 2024; Fattinger et al., 2019; Moreira et al., 2021; Ngo, Martinetz, Born, & Molle, 2013; Papalambros et al., 2017; Wunderlin et al., 2023; Zeller et al., 2024), to potentially alter neuropathological hallmarks and consequently delay or reverse AD symptoms.

Overall, we show direct and differential associations between SWA increase, plaque deposition, and memory function after a prolonged treatment of cohorts at different disease stages, suggesting sleep to be a promising therapeutic target for neuropathology and cognitive impairment in AD. Moreover, our finding that sleep has a restorative effect on cognition after early onset interventions proposes that identifying individuals at risk for developing AD before significant neuropathological progression is critical to arrest or delay the onset and/or progression of the disease.

## CRediT author statement

Sedef Kollarik wrote the initial manuscript; collected the data, performed the statistical analyses, and interpreted the data. Dorita Bimbiryte and Aakriti Sethi contributed to the collection and investigation of the data, reviewed and edited the manuscript; Inês Dias and Carlos G. Moreira contributed to the data analysis and curation; reviewed and edited the manuscript. Daniela Noain conceptualized, supported and supervised the study, and wrote, reviewed and edited the manuscript. All authors approved the final draft.

## Ethics approval statement

All experiments were approved by the veterinary office of the Canton Zurich and conducted according to the local and federal guidelines for care and use of laboratory animals under license ZH210/17.

## Acknowledgements

This work was supported by the Neuroscience Center Zurich (ZNZ) through the patronage of Rahn and Bodmer Co. (DN) and the Dementia Research - Synapsis Foundation Switzerland via an earmarked donation of the Armin & Jeannine Kurz Foundation (DN). The authors would like to thank Prof. Dr. med. Christian R. Baumann for his generous support.

